# Slow rhythmic eye motion predicts periodic alternation of bistable perception

**DOI:** 10.1101/2020.09.18.303198

**Authors:** Woochul Choi, Hyeonsu Lee, Se-Bum Paik

**Affiliations:** Institute of Information Electronics, Korea Advanced Institute of Science and Technology, Daejeon 34141, Republic of Korea; Department of Bio and Brain Engineering, Korea Advanced Institute of Science and Technology, Daejeon 34141, Republic of Korea

## Abstract

Bistable perception is characterized by periodic alternation between two different perceptual interpretations, the mechanism of which is poorly understood. Herein, we show that perceptual decisions in bistable perception are strongly correlated with slow rhythmic eye motion, the frequency of which varies across individuals. From eye gaze trajectory measurements during three types of bistable tasks, we found that each subject’s gaze position oscillates slowly(less than 1Hz), and that this frequency matches that of bistable perceptual alternation. Notably, the motion of the eye apparently moves in opposite directions before two opposite perceptual decisions, and this enables the prediction of the timing and direction of perceptual alternation from eye motion. We also found that the correlation between eye movement and a perceptual decision is maintained during variations of the alternation frequency by the intentional switching or retaining of perceived states. This result suggests that periodic bistable perception is phase-locked with rhythmic eye motion.

## Introduction

Bistable perception is a phenomenon by which subjects’ perceptions of ambiguous stimulus alternate spontaneously between two possible interpretations^1^. This perceptual alternation occurs periodically under various stimulus conditions associated with visual^2^ or other modalities^3^. In general, bistable perception is considered to be a consequence of the active interpretation of ambiguous sensory information, and it may provide important clues with which to understand the detailed mechanisms of sensory information processing in the brain.

The hallmark of bistable perception is the periodic alternation between two perceived states^4,5^. It has been observed that the average frequency of perceptual switching in an individual for a given stimulus (e.g., a Necker cube) is fairly consistent, while it largely varies across individuals ^4,6–9^. This raises questions about the mechanism of periodic perceptual alternation and about the determinants of the switching frequency in individuals. Regarding this issue, human psychophysics and neuroimaging experiments have demonstrated that various cognitive factors such as intention^10–13^, attention^14–18^ or adaptation^7,9,19–22^ can modulate the perceptual alternation frequency, and parameters of a neural basis, such as the grey matter volume^23–25^ and average inhibition level^20,26–29^, appear to be correlated with the frequency of perceptual alternation. For example, it was observed that asking subjects intentionally to switch or hold their perceptual states can significantly bias their perceptual alternation rate of bistable perception^10,11,13,30^. While such behavioral changes stemming from various factors have been observed, the detailed mechanism by which these components mediate perceptual alternation frequency remains unknown^18,31^.

While neuro-imaging studies have elucidated how neural activity patterns are correlated with alternations of perceptual decision^27,32–34^, other approaches have focused on eye movements, a crucial component of visual perception that may trigger measurable changes in various cognitive tasks. Studies of this issue have shown that for subjects who perceive illusory motion from a static image (e.g., a rotating snake and an enigma), micro-saccadic eye movements precede the active interpretation of the stimuli^35,36^, implying a causal relationship between eye movement and motion perception. Other studies also suggest that pupils tend to dilate before each perceptual decision during various types of bistable perception tasks^37,38^ (for a rebuttal, however, see ^39,40^). Another study illustrated how eye gaze positions while viewing a Necker cube may provide a negative feedback signal which suppresses the current status of visual perception^14^. Given these findings, we hypothesized that a precise estimation of eye movement can predict the profile of the perceptual alternation of a bistable stimulus.

To test our hypothesis, we simultaneously measured eye movements and perceptual behavior while human subjects performed tasks with three different types of stimuli. From this, we found that eye gaze positions oscillate slowly, with a frequency similar to that of individual perceptual alternation in bistable tasks. Furthermore, when we measured the average eye gaze positions before two possible perceptual decisions, we observed that the two eye gaze trajectories are in the opposite direction before two distinct perceptual decisions are made. Using a linear classifier of preceding eye gaze trajectories, we could predict subjects’ behavioral responses to ambiguous stimuli with approximately 90% accuracy. When the subjects were asked to switch or hold their perception intentionally, we found that the behavioral response was still predictable with gaze trajectories, despite the fact that the intentional control significantly alters both perceptual behavior and eye gaze oscillation. Taken together, our observations suggest that slow rhythmic eye gaze motion is tightly correlated with the periodic alternation of bistable perception, the frequency of which varies widely across subjects. This implies that oculomotor interaction is a crucial component when determining the subject-specific dynamics of perception under ambiguous circumstances.

## Results

### Rhythmic eye gaze oscillation is correlated with the periodic alternation of bistable perception

We undertook a human psychophysical experiment using three types of bistable stimuli: a Necker cube, a racetrack, and a rotating cylinder. Each subject’s eye gaze positions and pupil dilation (Fig. 1A) were recorded simultaneously. As observed previously, we confirmed that subjects’ perceptual responses to all three stimuli alternate periodically. The distribution of the phase duration – defined as the time spent on either of the two possible interpretations – was well fitted to a log-normal distribution. The peak value of this distribution was measured in each individual^4,41^. The eye gaze position during the task was also measured, and we found that an autocorrelation analysis of the 2-D positions of the eyes showed rhythmic oscillation (Fig. 1B), the frequency of which was estimated from the position of the first-order peak in the autocorrelation. In all three stimuli conditions, the oscillation period of the eye gaze and the period of bistable alternation appeared to be positively correlated (Fig. 1C, Pearson’s correlation r = 0.37, p < 1.6e-5, N = 132 measurements from all three experiments from 45 subjects). This result suggests that rhythmic eye motion during bistable perception is tightly correlated with the frequency of perceptual alternation in individuals.

**Figure 1.**
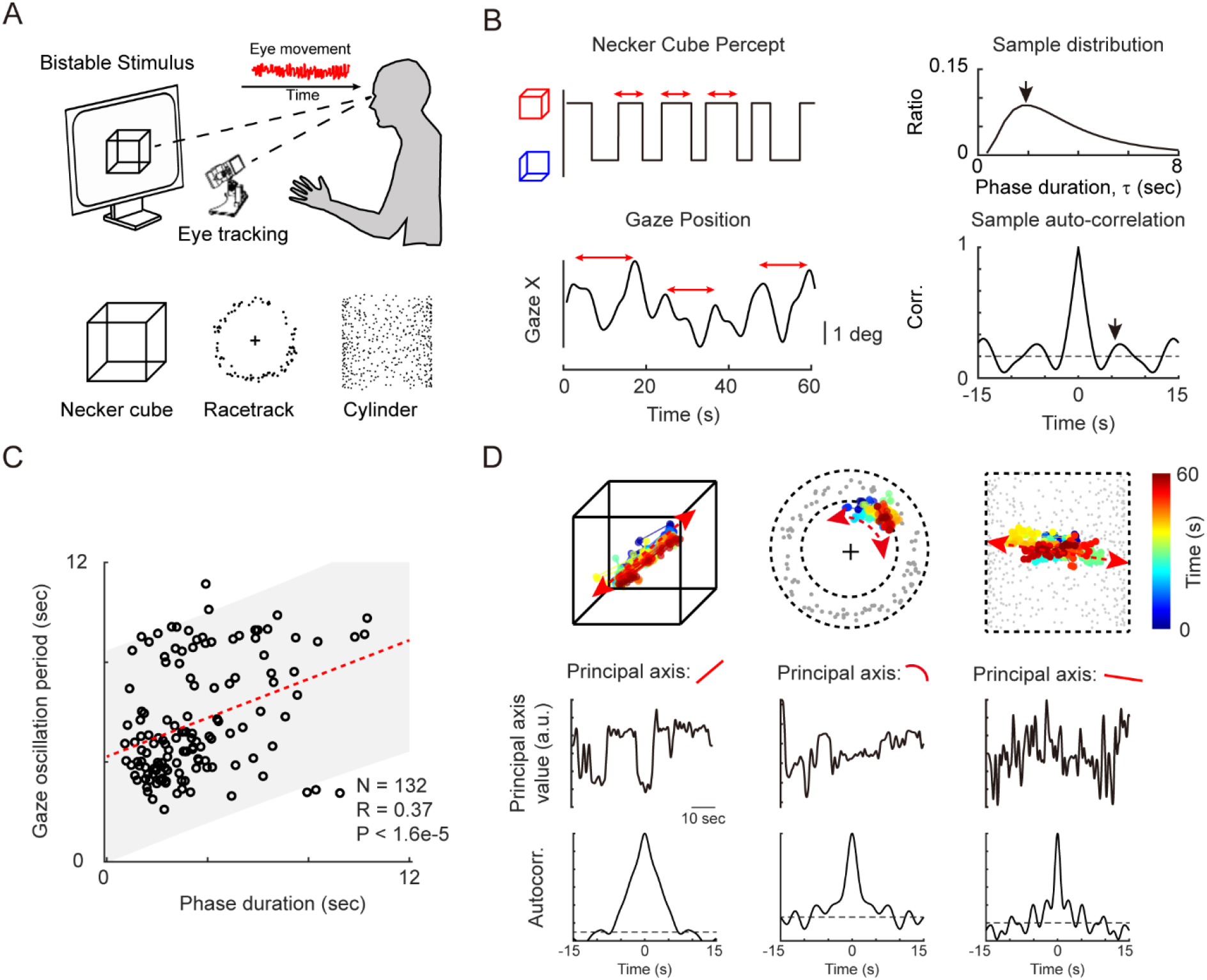
Rhythmic eye movement is correlated with the perceptual alternation of ambiguous bistable stimuli: (A) Simultaneous tracking of the eye gaze during three bistable stimuli. A Necker cube is a static stimulus and a racetrack and cylinder are moving stimuli consisting of black dots. (B) Sample perceptual response pattern and horizontal gaze position. In the perceptual response, the phase duration – time spent on each percept – was measured and the peak of the phase duration was denoted (top). From the gaze positions, the auto-correlation was calculated, and the first peak from the autocorrelation curve was denoted as the period of the gaze oscillation (bottom). (C) The gaze oscillation period and phase duration (from behavior) were positively correlated with each other (Pearson’s correlation, r=0.37, p< 1.6×10^−5^, N=132 measurements, collapsing all three experiments from 45 subjects). (D) Sample gaze positions and oscillation on the principal axis. Sample gaze positions for three stimuli are shown. The principal axis was calculated from binocular 2-D gaze positions. Note that the principal axis shows specific stimulus characteristics, such as a diagonal line for the Necker Cube, and the gaze oscillates on the principal axis (bottom).

We also analyzed the spatial profile of the rhythmic gaze motion in each stimulus condition. We found that the gaze positions are located on a specific axis in 2-D space in most conditions. For example, when the Necker cube stimulus is given, most subjects’ gaze positions were positioned along a diagonal line that links two opposite vertices of the cube (Fig. 1D). On the other hand, for the racetrack stimulus, gaze positions were aligned with the circumference, matching the perceived direction of illusory motion. For the 3-D cylinder stimulus, gaze positions appeared to be distributed on the vertical axis, on which the depth of the cylinder is perceived (Supplementary Figure 1). This result reveals that the rhythmic gaze motion oscillates along spatial axes that provide information with which to make perceptual decisions for each type of bistable stimulus condition.

### Distinct gaze trajectories before perceptual alternation in all three stimuli

Next, to examine the relationship between gaze motion and perceptual alternation further, we analyzed the gaze motion profile just before each perceptual response (bistable alternation of perception) is made (Fig. 2A); we measured the gaze velocity, saccade rate, micro-saccade rate, and pupil dilation before and after the onset of subjects’ responses of perceptual alternation. While we found no significant changes in the patterns of the gaze velocity, and the rate of saccadic/micro-saccadic motion around the subjects’ responses, we observed noticeable dilation of the pupil approximately 1s after the subject’s response (Supplementary Figure 2). Because the pupil dilation observed here occurred after perceptual decisions, it may be a reflection of certain parameters pertaining to each perceived state, rather than being related to the triggering of the perceptual response (see refs ^37,39,40^).

**Figure 2.**
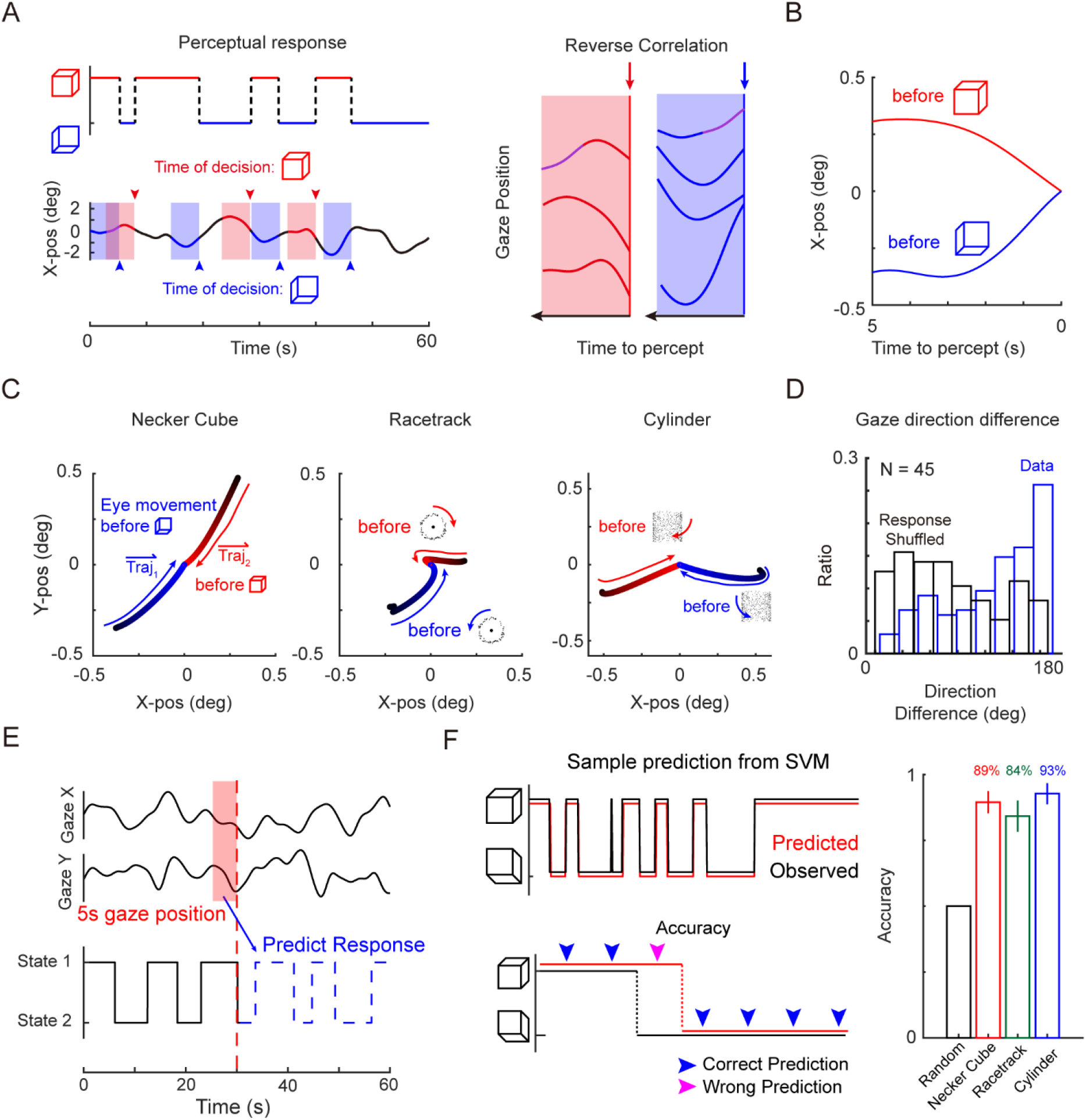
A slow rhythmic eye gaze predicts the perceptual decision of bistable perception: (A) (left) Horizontal eye movements during the perceptual alternation of ambiguous bistable perception. The shaded area denotes the five seconds before each perceptual decision. (right) Eye gaze position aligned with the perceptual decision. Reverse correlation calculates the average eye gaze position before each decision. (B) Sample reverse correlation result. Red and blue positions are the average gaze positions before each perceptual state. (C) Sample average eye gaze trajectory in two-dimensional visual space. For each subject and stimulus, gaze trajectories 1 and 2 are calculated and analyzed. (D) The gaze direction difference between trajectory 1 and trajectory 2 is calculated. The blue bar denotes the direction calculated from the data and the black bar denotes the direction calculated from the shuffled behavior response. (E) Predicting the perceptual response from the gaze position using a support vector machine. Horizontal gaze positions, vertical gaze positions, and derivation of the gaze positions of five seconds were used to predict the perceptual responses of individuals. (F) Prediction accuracy of the SVM. A sample prediction of the perceptual responses, and the observed responses are shown in red and black (left). For all three stimuli, the prediction accuracy exceeded 80% (right).

More importantly, we observed distinct eye gaze patterns, approximately within a 5s window before each perceptual alternation (Fig. 2B). In all three stimulus conditions, the average gaze trajectories before each perceptual switch were in the opposite direction in 2-D space. Sample average trajectories from three different stimuli are shown in Fig. 2C. While the trajectories of the two eyes deviated largely across individuals, we confirmed that the average of two trajectories were in opposite directions consistently in all three stimuli conditions and for all individuals tested (Fig. 2D). This result reveals a stereotypic eye gaze pattern before each bistable alternation, the spatial axis of which is aligned to the principal axis of the gaze trajectories during bistable tasks, as indicated in Fig. 1D.

These observations imply that rhythmic eye movement changes accompany, or even may trigger, each perceptual alternation. To test whether this distinct gaze motion contains enough information to predict the future timing of perceptual alternation, we trained a linear support vector machine (SVM) to predict the perceptual status of the subsequent frame in individuals using only the preceded gaze position information (Fig. 2E). We found that the future perceptual status in all three stimuli can be predicted with up to 90% accuracy from gaze positions within a 5s window previously (Fig. 2F, Supplementary Figure 3). These results suggest that the profile of slow rhythmic eye motion contains enough information to estimate individuals’ perceived status with ambiguous visual stimulus.

### Correlation between gaze motion and bistable alternation is maintained under top-down control

Next, we investigated whether the observed correlation between perceptual alternation and gaze motion can be altered by top-down control. Previous studies reported that the periodic switching of bistable perception can be significantly modulated by intentional control^10–12^. Considering this, we investigated changes of the correlation between perceptual alternation and gaze motion while the frequency of bistable alternation is altered by intention: we measured the perceptual responses and gaze motions of each subject with 1) no intention, 2) an intention to switch, and 3) an intention to hold their perceived status (Fig. 3A). As expected, first we found that subjects’ frequencies of perceptual alternation changed as they intended to switch or hold their current state (Fig. 3B). Intriguingly, the gaze motions also appeared to be altered significantly with this intentional control (Fig. 3C) such that the frequency of eye motion increased or decreased similarly to the changes of perceptual alternation. The phase duration of the perceived status in all stimuli increased/decreased significantly when the subjects intended to hold/switch their perception (Fig. 3D, Freidman test, p< 1.0e-4 for all three stimuli, N=23). Similarly, the oscillation period of the eye gaze was increased/decreased significantly by top-down intention (Fig. 3E and E, Friedman test, p<0.0017, p<0.019, p<0.0316, for the Necker cube, racetrack, and rotating cylinder, respectively). Overall, both the perceptual behavior and the gaze motion were noticeably altered when the subjects intended to change their behavior (Fig. 3F).

**Figure 3.**
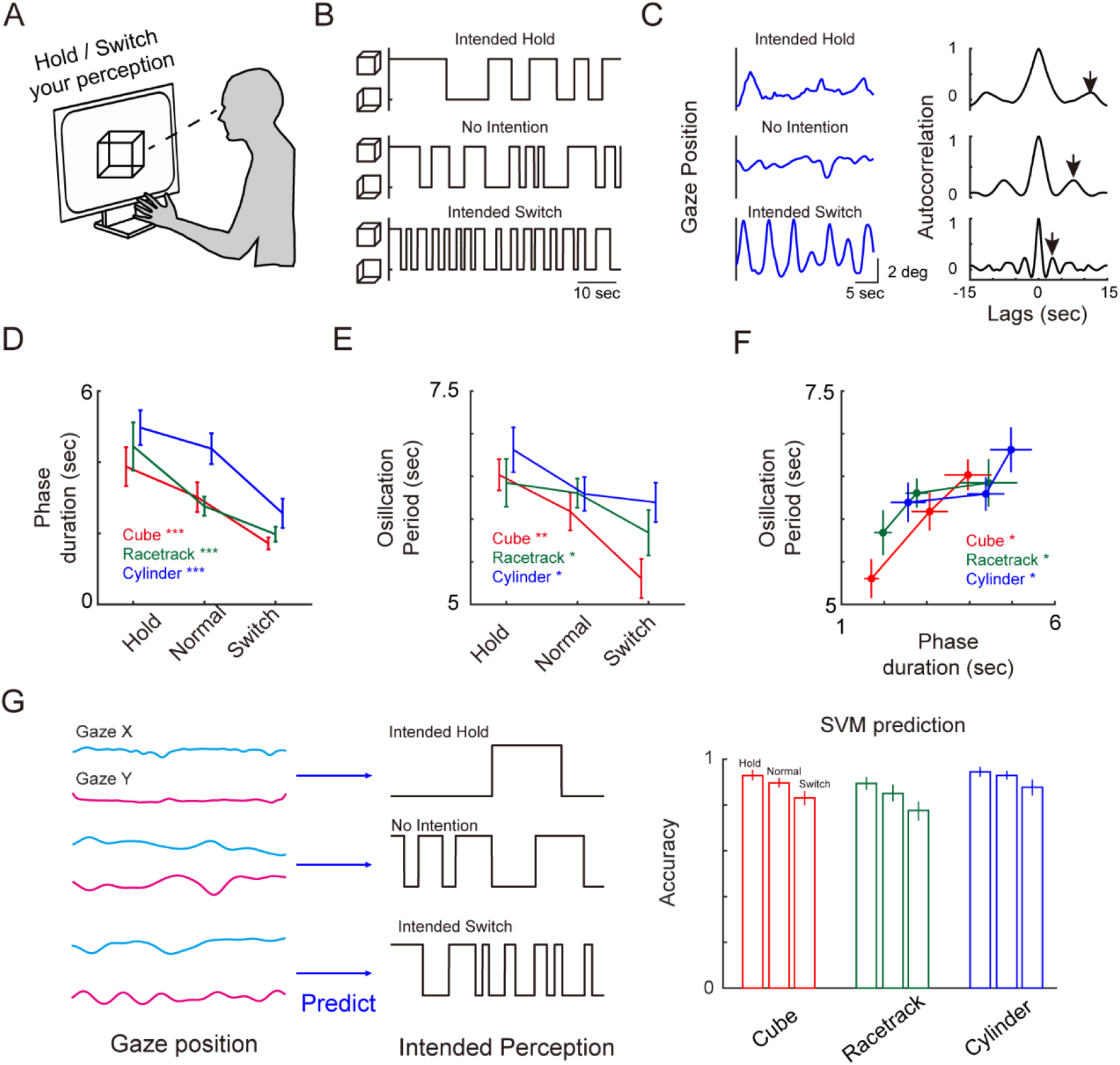
Intentional control alters both perceptual behavior and eye movements, while the relationship between eye movement and perception remains intact: (A) Subjects were instructed to hold/switch their perceived states as long/rapidly as they could. (B) Sample perceptual behavior under intentional switch/hold. (C) Sample eye gaze trajectories (left) and their autocorrelation (right) under intentional switch/hold. The first positive peak from the autocorrelation curve is shown. (D) Altered phase duration under intentional switch/hold conditions. The phase duration was significantly altered as subjects intended to switch/hold their perceptual states (Freidman test, p< 1.0^−4^ for all three stimuli, N=23 for each stimulus). (E) Altered eye gaze oscillation under intentional control. Similar to the phase duration, the gaze oscillation period was significantly altered for all three stimuli (Friedman test, p<0.0017, p<0.019, p<0.0316, for the Necker cube, racetrack, and rotating cylinder, respectively) (F) Correlation between the phase duration and the gaze oscillation under intentional control. The average phase duration and oscillation period for each stimulus are shown. (G) Prediction accuracy of the SVM. Similar to Figures 2E and F, the perceptual responses were predicted from the preceding eye gaze positions. For all three stimuli and in all intended conditions, the prediction accuracy exceeded 75%. Note that in all three conditions, a slower perceptual alternation (hold) was easier to predict than a faster perceptual alternation (switch).

However, we observed that the relationship between perceptual alternation and gaze motion remained under intentional control. Using the same procedure shown in Fig. 2E, we found that each subject’s perceptual responses could be well predicted by information of the gaze positions within a 5s window previously (Fig. 3G), despite the fact that both the eye gaze and perceptual alternation behavior were significantly altered. This suggests that the relationship between gaze motion and perceptual alternation is intact under top-down intentional modulation.

### Eye motion during free-viewing is correlated with the temporal dynamics of perceptual alternation

Lastly, we investigated the possibility that the profile of eye gaze is correlated with the frequency of perceptual alternation, not only in certain bistable conditions but also in a normal condition such as free-viewing, as eye motions during active perceptual tasks involve a number of untraceable components, such as intention, the attention level, action planning, and anticipation. To examine whether the temporal frequency of “normal” eye motion is correlated with the subject-specific frequency of bistable perception, we measured eye motions during free-viewing, i.e., with no cognitive task required. To do this, we asked the subjects to observe freely 100 natural images, for three seconds per image^42^. During this free-viewing task, the frequency of micro-saccade events, fixation events, and fixation durations were measured. Intriguingly, subjects with more frequency alternations during bistable perception tended to explore images more actively (more spatial navigation across locations) compared to subjects with less frequent alternations in bistable tasks (Figs. 4A and B). Thus, the frequency of fixation event during the free-viewing of a natural image is positively correlated with the frequency of alternation in bistable perception (Fig. 4C, left, Pearson’s correlation r = 0.50, P<0.006). Similarly, the fixation duration in free-viewing is negatively correlated to the frequency of bistable alternation (Fig. 4C, middle, Pearson’s correlation r = −0.52, p<0.004, N = 29). Furthermore, the frequency of micro-saccade events also appeared to be correlated positively with the frequency of alternations in bistable perception (Fig. 4C, right, Pearson’s correlation r = 0.63, P<0.0002). These results imply that eye motions observed not only during cognitive tasks but also during natural conditions reflect the intrinsic temporal dynamics of the perceptual process in individuals.

**Figure 4.**
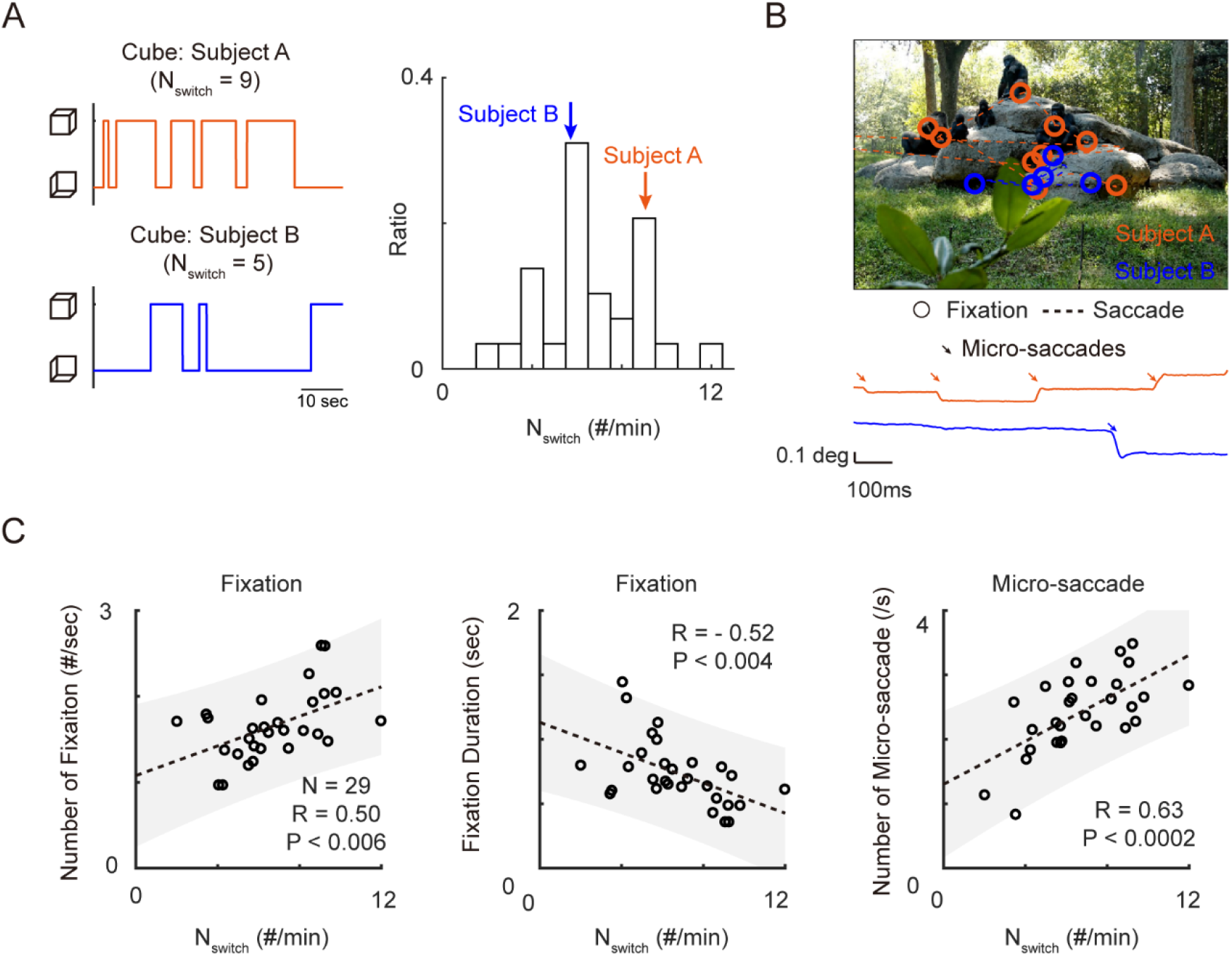
Saccadic eye movement during free-viewing is strongly correlated with the temporal dynamics of a perceptual alternation: (A) Two sample subjects with comparative perceptual dynamics. Subject A has a high number of perceptual alternations (N_switch_ = 10.2), while subject B has a relatively low number of perceptual alternations during 60 seconds with the Necker cube trial (N_switch_ = 4.7) (B) Sample eye gaze trajectories during the free-viewing of natural images. The circle denotes the fixation and the dashed line denotes the saccadic traces (top). Micro-saccade traces are denoted as arrows during the free-viewing task (bottom). Note that subject A explores more spatial locations compared to subject B during three seconds of free-viewing. (C) Correlation between saccadic eye movements and the number of perceptual alternations. The average number of fixations was positively correlated with N_switch_ (left, Pearson’s correlation r = 0.50, P<0.006, N=29), and the average fixation duration was negatively correlated with N_switch_ (middle, r = −0.52, p<0.004, N = 29). The average frequency of micro-saccade events was positively correlated with N_switch_ (right, r = 0.63, P<0.0002, N = 29).

## Discussion

The main finding of the current study can be summarized as follows. The temporal dynamics of eye motion is tightly correlated with the temporal alternation of active perceptual behavior in individuals. During the bistable perception task, we found that the subjects’ perceptual behavior and gaze motion both oscillate at a slow, idiosyncratic frequency that varies across subjects. Notably, gaze motions are aligned with a specific axis in space, along which the status of an ambiguous stimulus can be perceived differently. We found that preceding eye motions can accurately predict the future perceptual status of individuals. We also found that this relationship between eye gaze and perceptual alternation is maintained even when the eye motion and perceptual response are both strongly biased by intentional control. Lastly, we found that saccadic eye motion with no cognitive demand – i.e., a free-viewing task – is also correlated with the temporal alternation of perceived status in individuals. These findings suggest that the correlation between the eye motion and perceptual behavior is tightly correlated.

Inter-individual variability of the perceptual alternation frequency is considered to provide information about visual processing in each individual^25,41,43–46^ and about how the same stimulus is differently encoded in different brains^47–49^. The phase duration (or inversely, alternation rate) in bistable perception appears to vary widely across subjects and is thought to reflect the distinct processing time of individuals to perceive information in the external world^5,41^. The slow, rhythmic eye motion observed in the current study is correlated with this perceptual alternation frequency, implying that this eye motion also provides information about visual processing in individuals. One notable feature of this eye motion is that it is slower than other characteristic eye motions previously observed. While the eyes can move on various time scales – from very fast tremors^50^ or drift^51^ to relatively slow micro-saccades (frequency typically varying around 1Hz^52–55^) – the eye oscillation observed here has a period varying from 0.25 to 1Hz, as slow as the eye movement during REM sleep^56^. This finding reveals a new type of eye dynamics that must be considered as important when attempting to understand the perceptual behaviors of human subjects during active perceptual tasks.

Another interesting point is that this gaze motion is observed with both stationary (Necker Cube^14,57^) and dynamic stimuli involving motion (racetrack and rotating cylinder). Furthermore, this gaze motion appears to be aligned with a specific axis of the bistable stimuli, in which the perceived status of the stimuli can be altered. For example, for the Necker cube, most subjects’ gaze positions were biased along the line linking two opposite vertices of the cube. As it was previously reported that the gaze position is a crucial factor related to the switching of the perceived status^57,58^, our finding also provides information about a more general relationship between gaze motion and perceptual alternation.

Combined with the fact that 1) the eye gaze oscillates with a similar period of perceptual alternation (Figs. 1C) and 2), the two gaze trajectories before each perceptual decision had opposite directions (Figs. 2A-D), the preceding eye gaze positions would contain enough information to predict the future perceptual responses. As expected and confirmed by the linear classifier, the preceding eye gaze positions can accurately predict the subject’s future perceptual interpretation (Figs. 2E and F). This is consistent with a previous study that showed that the horizontal optokinetic response (OKN) can predict perceptual responses while subjects observe a bistable plaid motion stimulus^59^, but our study shows that eye gaze dynamics not only predicts a dynamic bistable stimulus but can also be used to predicts a stationary bistable stimulus – a Necker cube in this case. This clearly shows that the preceding eye gaze positions are largely related to how individuals will interpret an ambiguous visual stimulus. Note that a causal relationship between eye movement and perceptual alternation is difficult to reveal due to the nature of the task, in which the time point of perception (i.e., when the brain interprets a rotating dot in the clockwise direction) and the motor response (i.e., when a finger touches a keyboard that corresponds to clockwise rotation) are intertwined with each other^33,34^. Although we could not directly show a causal relationship indicating that eye movement drives perceptual alternation, from the fact that the eye positions even three to five seconds before the two decisions were significantly opposite (Figs. 2B and C), we speculate that the eye gaze clearly contains perceptual information regardless of the motor response or action planning process.

Several possible scenarios can explain the observed correlation between eye motion and perceptual alternation: 1) The brain undertakes perceptual alternation without involving eye movements, but the eye moves in a specific direction as a consequence of the perceptual response. 2) The eye moves in a certain direction which drives perceptual alternation. 3) A third factor determines both the eye movement and perceptual alternation in cases of ambiguous visual stimuli. While our result cannot provide a definite answer, we suggest a combination of the second and third scenarios to explain the mechanism of perceptual alternation. We speculate that the first hypothesis could be rejected because the eye positions and gaze trajectories emerge very early before the perceptual decision is made. As Einhauser and colleagues suggested^14^, we propose that when eye positions approach certain extreme points in ambiguous stimuli, the percept becomes more likely to alternate from the current percept, a contention supported by our reverse correlation result (Figs. 2A-C). Moreover, the relationship between the eye gaze and perceptual alternation was largely maintained under intentional control, suggesting that eye gaze and perceptual alternation are both governed by common factors (Fig. 3).

We subsequently examined whether there remains a relationship between eye movement and perceptual alternation dynamics even if there is no cognitive process during the task – i.e., action planning, motor response, attention, and intention, for instance. Surprisingly, the characteristics of eye gaze patterns during free viewing were also significantly correlated with individual perceptual temporal dynamics. The frequencies of saccade and micro-saccade events were positively correlated with the phase durations of individuals (Figs. 4B and C). It is also important to note that micro-saccades cannot be controlled by one’s will, suggesting that the saccadic frequency is an intrinsic characteristic of individuals. Combined with a previous report positing that the phase duration of the bistable perception reveals the time window of individuals for integration visual information^41^, this result suggests that the eye dynamics reflect the intrinsic visual information integration process. For example, individuals with long visual integration times may move their eyes slowly, with fewer explorative saccades, while individuals with short visual integration times may move their eyes rapidly, making long explorative saccades. These results collectively suggest that the intrinsic characteristics of individuals determine the eye gaze patterns and perceptual alternation dynamics.

This leads to the question of how the relationship between eye movement and active perceptual process can help us understand the visual circuits involved. Computational model studies suggest that diverse visual functions – from the fundamental orientation selective response^60–62^ to complex cognitive behaviors such as face perception or working memory^48,63–65^ – can arise from a simple variation of the physical circuit structure. If this is the case in the present study, we can also suggest that a common neural circuit mechanism with simple variation determines both eye movement and active visual perception from the intact correlation between intrinsic eye movement and the perceptual alternation dynamics^66–69^. While it is still controversial as to which neural activity drives the interpretation of ambiguous stimuli^16,23,25,28,32–34,41,70–74^, various computational models and human experiments suggest that the inhibition or adaptation level determines the temporal dynamics of perceptual decisions^9,13,26,27^. In terms of the neural mechanisms of eye movements, various brain areas – including the frontal eye field, the cerebellum, and the superior colliculus – are involved in the eye movement process. Specifically, inhibitory signals from the cerebellum or superior colliculus are largely involved in eye movement control. Taken together with the finding that saccadic eye movements during free-viewing also reflect inhibition level^75–77^, a high inhibition level in the cerebellum or superior colliculus may induce in slow perceptual alternation during bistable perception and slow saccadic (rhythmic) eye movements. Further studies pertaining to inhibitory levels in such visual circuits may reveal the overall mechanism in terms of eye movements and active visual perceptual processes.

To summarize, we demonstrated that intrinsic and distinct rhythmic eye movements precede the active interpretation of various types of ambiguous stimuli. These rhythmic eye gaze oscillations well predict individual future perceptual decisions, and the rhythmic oscillations and predictions were retained when subjects controlled their intention. We have showed that eye movements without any cognitive tasks were also correlated with the temporal dynamics of a perceptual decision, suggesting that an intrinsic factor of the common neural circuit may determine both eye movement and active perception.

Taken together, we propose that rhythmic eye movements reveal individual active visual interpretation processes of ambiguous visual stimuli.

## Methods

### Participants

Forty-five subjects (with normal or corrected normal vision) were enrolled in the experiment. All experimental procedures were approved by the Institutional Review Board (IRB) of KAIST and all procedures were carried out in accordance with approved guidelines. Written informed consent was obtained from all subjects.

### Display and visual stimulus

Visual stimuli were presented on an LCD monitor screen (DELL U3014, 29.8 inches, 2560 × 1600, 60 Hz temporal resolution) for all experiments. Subjects were positioned 120 cm away from the monitor and were asked to report their perception of each stimulus using buttons on the keyboard. For active perception tasks, three bistable stimuli were used: a Necker cube, a racetrack, and a rotating cylinder. For the Necker cube task, subjects reported a simultaneous perceptual state of Necker cube formation with two keys on the keyboard. For the racetrack task, subjects reported the direction of rotating dots – clockwise or counter-clockwise, or mixed rotation. For the rotating 3D cylinder task, each subject was asked to report the direction of the rotating cylinder – clockwise or counter-clockwise (viewed from the top), or mixed rotation.

The Necker cube stimulus consisted of 12 identical lines, each of which had a length of 5.5 degrees and a width of a 1-minute degree. The racetrack model stimulus consisted of randomly distributed black dots in the annulus area. The inner and outer radii of the stimulus were 3.5 degrees and 5 degrees, respectively, from the center of the screen. The movie frame was refreshed every 50 ms, generating a different distribution of dots. The size of an individual dot was 5 min in diameter, and the density of the dots was set to 5 dots/deg^2^. To create a fixation point on the screen, a black cross was positioned at the center of the screen, and each subject was asked to fix their eyes on the cross during the racetrack experiment. The rotating 3D cylinder stimulus consisted of black dots with a density of 3 dots/deg^2^. First, randomly distributed dot positions were plotted on the surface of a cylinder (7-degree diameter and 7-degree height) and then projected onto a 2D screen with a 7-degree tilted view angle from the top. Movie frames were refreshed every 33 ms, and the rotation speed was set to 4 degrees/frame. The stimulus conditions were optimized based on the results from preliminary trials.

For the free-viewing task, 100 randomly selected images from the previous dataset were used^42^. Face images were intentionally excluded because the fixation of face images is known to concentrate in the areas of the eyes, nose, and mouth. Subjects viewed each image at a full resolution for 3 seconds separated by 1.5 seconds of viewing a gray screen with a fixation point in the center. Subjects were asked freely to explore the presented images without any intention. All visual stimuli were presented with MATLAB Psychtoolbox 3.0.

### Behavior

On the first day, subjects viewed the Necker cube, the racetrack on the second day, and the rotating cylinder in the last (third) day. On each day, the experiment was divided into two sessions; the first session was an intentional control session and the latter session was a moving fixation dot session. In the first session, subjects were trained to 1) normally respond to two interpretations of the stimulus, 2) intentionally switch their perceived interpretation as rapidly as they could, and 3) intentionally hold their current perceived interpretation for as long as they could. Each intentional condition was indicated before the beginning of the 60s task. To minimize confusion, two blocks of 24 intentional tasks consisted of consecutive 8 normal conditions, 8 switch condition and 8 hold condition (for a total of 48 trials – two blocks) were executed. The binocular eye was tracked during this session.

### Eye-tracking

Through the experiment, the binocular gaze position and pupil size were recorded using EyeLink 1000 Plus eye-tracker (SR Research) with a sampling rate of 500Hz. Subjects’ heads were stabilized with a chin-rest to minimize head movements. Before each experimental setup, subjects underwent a calibration session, with fixation points presented on the screen. The dominant eye was identified by the Miles test.

### Analysis

#### Eye gaze position

To extract a subject’s binocular 2-D eye gaze position during the task, 500Hz binocular eye gaze positions were first downsampled into 250Hz. The time windows before and after 100ms when the pupil disappears and reappears were considered as blink events. Subsequently, all eye gaze positions during a blink event or outside the monitor were nullified and were linearly extrapolated according to the neighboring eye gaze position. Next, the eye gaze position was smoothed with a 4ms Gaussian filter.

### Detection of saccades and micro-saccades

The detection of saccadic events and fixation events followed earlier research^42^. To illustrate the process briefly, if the eye gaze trajectory starts and ends with more than 6(deg/sec^2^) of acceleration, the trajectory was defined as a saccade. A trajectory below the acceleration threshold which lasts longer than 50ms was defined as a fixation event.

Detection of micro-saccades also follows earlier work^36^. To illustrate the process briefly, the event with the highest velocity at five events per second was deemed a micro-saccade candidate. Next, the peak velocity, initial acceleration peak, and final acceleration peak distribution were used as data points, and the k-means clustering algorithm was used to define the micro-saccade event that showed the largest average magnitude.

### Reverse correlation: Eye gaze position

To extract a subjects’ response-triggered eye gaze pattern, we initially measured the time point at which the perception changes from state 1 to state 2, t_1to2_, or from state 2 to state 1, t_2to1_. For every 60s trials, we extracted the two-dimensional binocular eye gaze position 15 seconds before every i^th^ response of a perceptual change time, t^i^_1to2_, or t^i^_2to1_, relative to eye gaze position at the time of perceptual change t^i^:

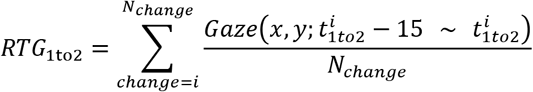

### Eye gaze oscillation

To extract oscillations from the 60-second-long eye gaze pattern, first we converted the 2-D gaze position into the distance from the center of the gaze. Next, we calculated the auto-correlation from the distance smoothed with a Gaussian filter, of which the width varies from 250ms to 1.5 seconds. The filter width which generates the highest first peak value was then chosen from the autocorrelation curve. From the autocorrelation curve of the selected width, the first peak was identified as an eye gaze oscillation indicator, and the peak position was defined as the gaze oscillation period. Detection of the peaks was computed with the MATLAB function ‘findpeaks’.

### Predicting perceptual responses from the eye gaze position

To predict the perceptual response from the eye gaze position, a linear support vector machine (SVM) was utilized^78^. To illustrate the process briefly, horizontal and vertical eye gaze positions were sampled with a 100ms interval. As the input to the SVM model, the raw x, y gaze position, the x, y gaze velocity, and the time-shifted gaze position and velocity were calculated at each time point. The SVM model was used to predict the next frame’s perceptual response, with ‘1’ and ‘-1’ labels for each perception. Ten-fold validation was performed, as the training utilized randomly selected time points from 60s. The gaze duration used for the prediction varied from 0.1 seconds to 8 seconds.

## Acknowledgments

This work was supported by grants from the National Research Foundation of Korea (NRF) funded by the Korean government (MSIT) (No. NRF-2019R1A2C4069863, NRF-2019M3E5D2A01058328) (to S.P.).

## Author contributions

S.P. conceived of the project. W.C. and S.P. designed the psychophysics experiments. W.C. performed the experiments. S.P. directed the experiments. W.C., H.L. and S.P. analyzed the data. W.C. and S.P. wrote the manuscript.

## Supplementary Information

**Supplementary Figure S1.**
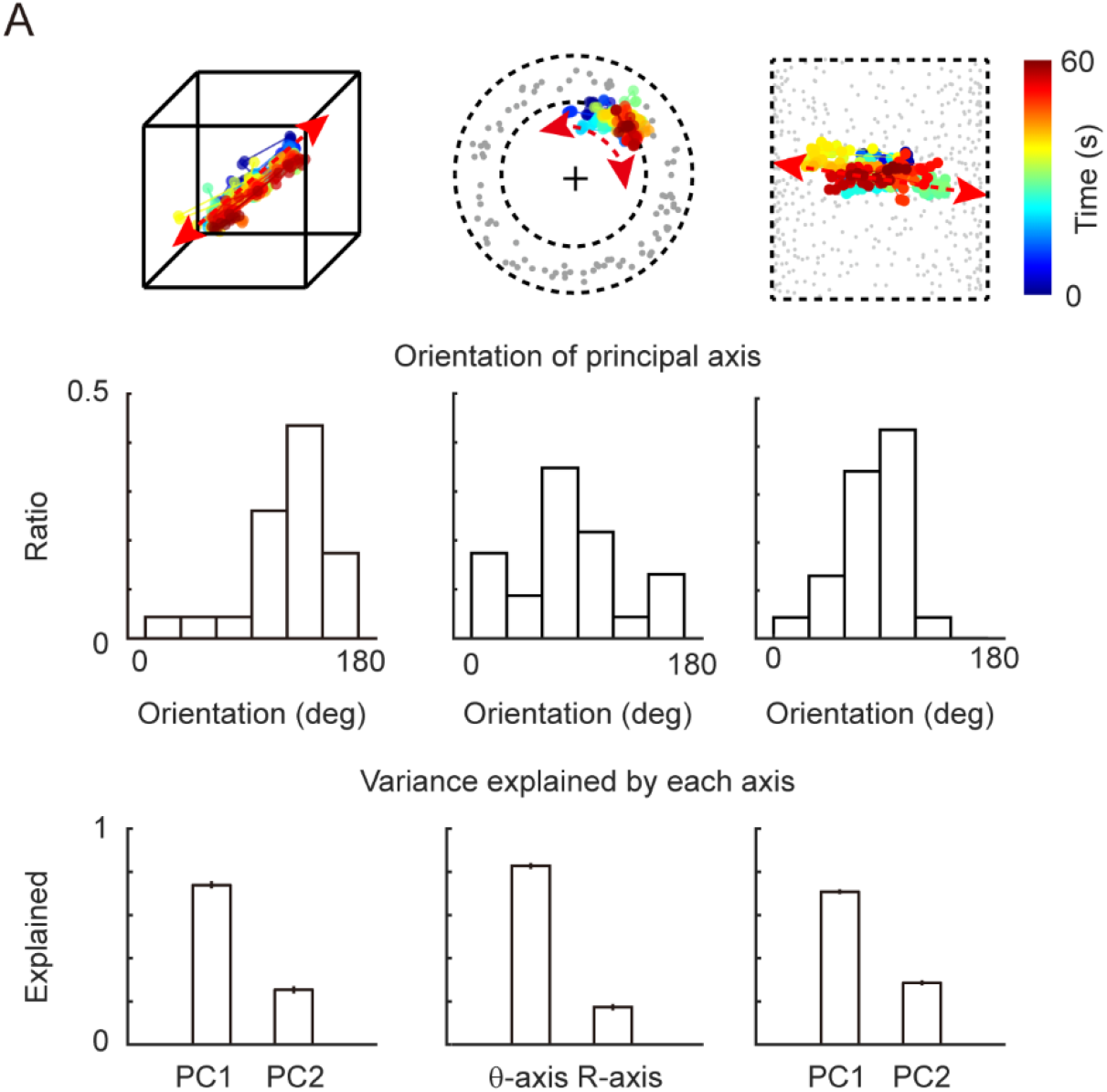
The principal component of gaze positions during three bistable perceptions. The top row shows the sample gaze positions during the three tasks. The middle row shows each individual’s principal axis direction for three different stimuli. The bottom row shows the explained variance percentage from the first principal axis and the second principal axis. Note that the co-variance was calculated on the r-theta plane in the racetrack condition.

**Supplementary Figure S2.**
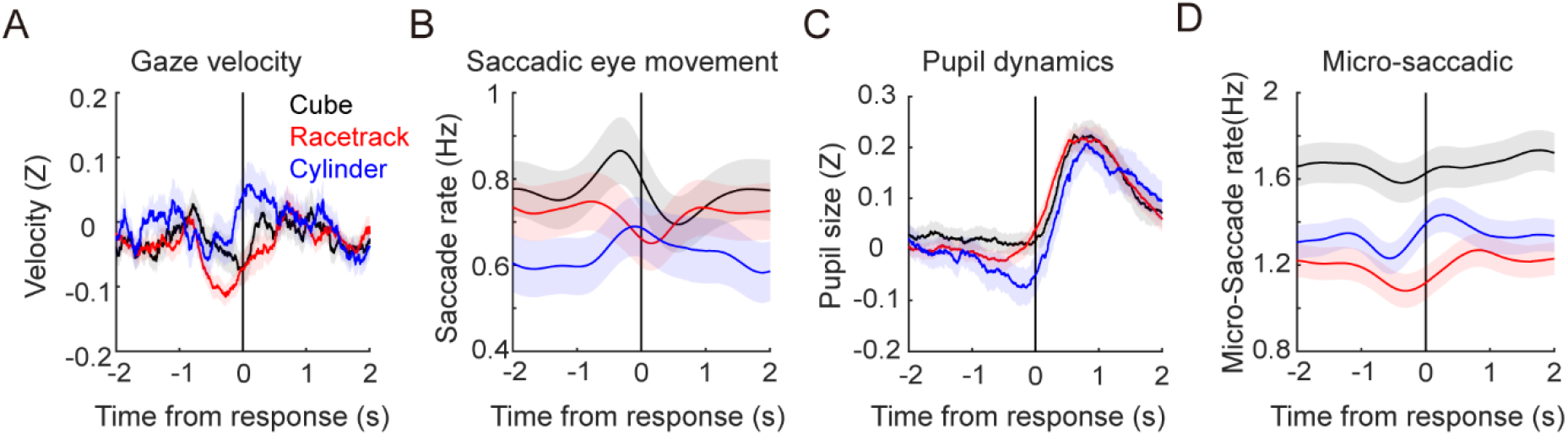
Reverse correlation of A) gaze velocity, B) saccadic eye movement rate, C) pupil dynamics and D) micro-saccadic eye movement rate before and after the perceptual decision. Note that the pupils dilate after the subjects make their decisions. Other parameters, in this case the gaze velocity, saccadic eye movement, or micro-saccadic movement rate, showed no significant trend before and after the perceptual response.

**Supplementary Figure S3.**
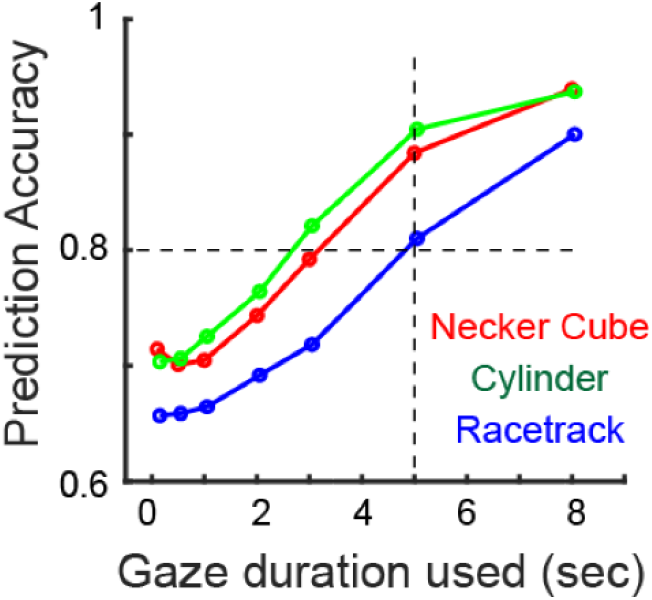
The perceptual state of ambiguous stimuli is predicted from the preceding eye gaze positions using a SVM (support vector machine). As more gaze positions were used to predict the perceptual response, the prediction accuracy increases. We chose a period of five seconds for gaze positions, resulting in >80% accuracy for all three stimuli.

**Supplementary Figure S4.**
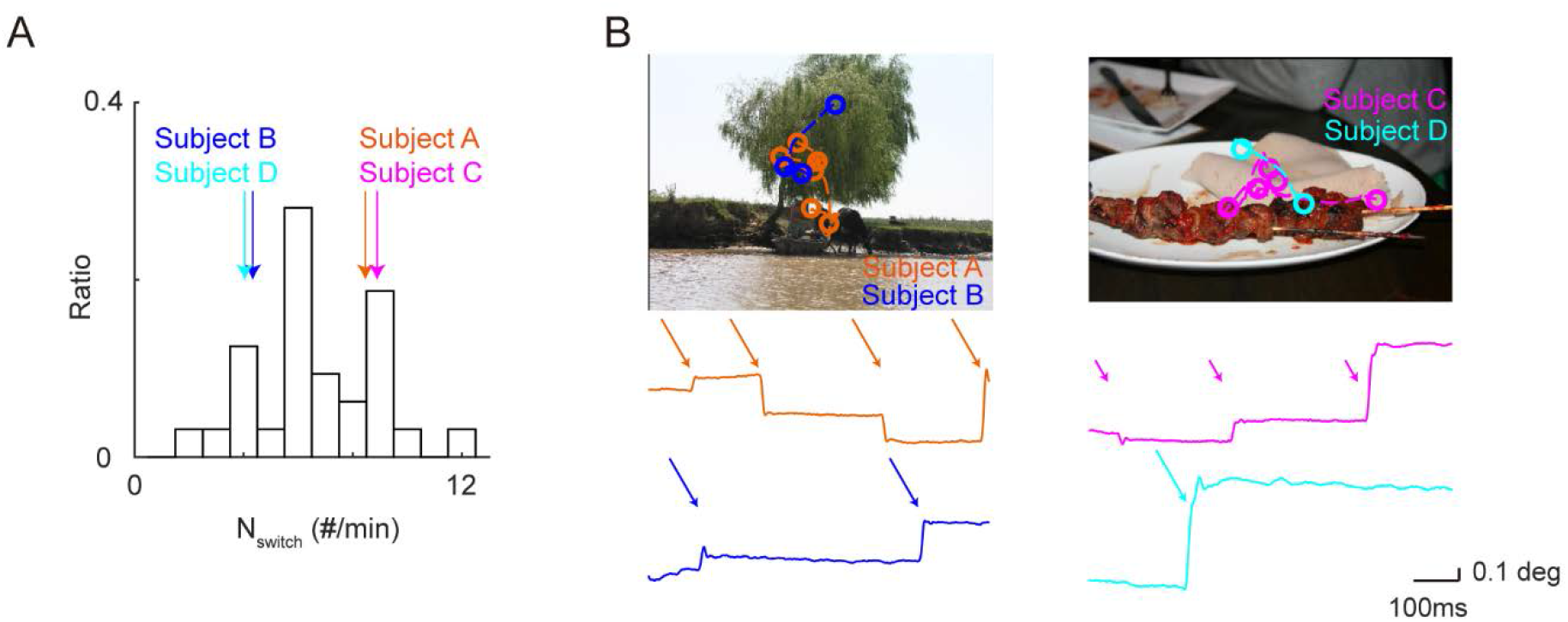
Fixation and saccadic eye movements during the free-viewing of natural scenes. Similar to Figure 4, subject C (i.e., fast alternations during bistable perception) makes more exploratory saccadic eye movements than subject D (i.e., slow alternation during bistable perception).

